# Haploid mouse germ cell precursors from embryonic stem cells reveal *Xist* activation from a single X chromosome

**DOI:** 10.1101/2020.10.30.361915

**Authors:** Eishi Aizawa, Corinne Kaufmann, Sarah Sting, Remo Freimann, Anton Wutz

## Abstract

Mammalian haploid cells have applications for genetic screening and substituting gametic genomes. Here we characterize a culture system for obtaining haploid primordial germ cell-like cells (PCGLCs) from haploid mouse embryonic stem cells (ESCs). We find that a haploid genome is maintained in PGCLCs with a high frequency indicating a substantially lower rate of diploidization than somatic cells. Characterization of the differentiating haploid ESCs reveals that *Xist* is activated from the single X chromosome. This observation suggests that X chromosome inactivation is initiated in haploid cells consistent with a model where autosomal blocking factors set a threshold for X-linked activators. The germline segregates from the epiblast and differs from somatic lineages in gene expression and epigenetic mechanisms. The ability of primordial germ cells for repressing *Xist* might contribute to the maintenance of a haploid genome.

## Introduction

In mice, the germline derives from cells of the proximal epiblast and primordial germ cells (PGCs) segregate to an extraembryonic location adjacent to the posterior end of the embryo. PGC differentiation has been recapitulated in cultures of mouse epiblast cells (Ohinata et al., 2009). Furthermore, functional gametes have been obtained from primordial germ cell-like cells (PGCLCs) derived from embryonic stem cells (ESCs) in mice (Hayashi et al., 2012; Hayashi et al., 2011). Progress in culture techniques is opening opportunities for studies of the mammalian germline (Hayashi and Saitou, 2013). Germ cells differ from the somatic lineages in several key aspects including gene regulation and epigenetic modification (Hill et al., 2018; Leitch et al., 2013). During migration to the genital ridges, erasure of genomic imprinting occurs in PGCs, and the inactive X chromosome (Xi) is reactivated in female PGCs (Sugimoto and Abe, 2007).

Mammalian dosage compensation is facilitated by inactivation of one of the two X chromosomes in female cells (Lyon, 1961). Therefore, a single transcriptionally active X chromosome (Xa) faces two sets of autosomes in the diploid genome. In mice, X chromosome inactivation (XCI) is initiated by the long noncoding *Xist* RNA, which is expressed from, and accumulates over the inactive X chromosome (Xi) before X-linked gene repression (Galupa and Heard, 2018). *Xist* is first expressed in four-cell stage blastomers and initiates imprinted XCI of the paternally inherited X chromosome (Okamoto et al., 2004; Takagi and Sasaki, 1975). Cells that are destined to form the embryo reactivate the Xi at the blastocyst stage. In the developing female epiblast two Xa are present before embryonic day (E) 5.5, when random XCI is initiated (Mak et al., 2004). Thereafter, the Xi is maintained in all somatic lineages, but in the female germline Xi reactivation is observed. From E7.0 two Xa are present in PGCs (Ohhata and Wutz, 2013; Sugimoto and Abe, 2007). These observations show the dynamics of the dosage compensation system.

The regulation of XCI remains to be fully understood. A number of observations suggest that the X to autosome (X:A) ratio controls the initiation of XCI and *Xist* expression. One model posits that autosomal blocking factors prevent *Xist* activation for explaining the observation that a single X chromosome is insufficient for initiation of XCI in male cells (Barakat et al., 2014; Kay et al., 1993; Pollex and Heard, 2019). In female cells, twice the number of X-linked activators overcomes an activation threshold for *Xist* leading to the initiation of XCI. Diffusible X-linked activators and autosomal blocking factors are beginning to be understood (Barakat et al., 2014; Pollex and Heard, 2019). An alternative model is based on the observation of paring of the X chromosomes at the X chromosome inactivation center (*Xic*) (Bacher et al., 2006; Xu et al., 2007; Xu et al., 2006). The *Xic* encompasses the *Xist* gene and other regulators of XCI including the antisense *Tsix* transcript. Genetic elements that are required for *Xic* pairing have been identified and shown to induce XCI when transgenically integrated into autosomes in male ESCs (Augui et al., 2007). Furthermore, stochastic regulation of *Xist* has been proposed from studies of tetraploid ESCs, where a variable number of X chromosomes displayed activation of *Xist* upon entry into differentiation (Monkhorst et al., 2008). Investigating the X counting mechanism in the context of different autosomal dosage has led to further understanding of the underlying regulation.

The mechanism of XCI has been extensively studied in ESCs, as these recapitulate the relevant developmental stages that would be difficult to access in the post-implantation embryo. In mice, female ESCs initiate XCI at the entry into differentiation. With the establishment of haploid ESCs (haESCs) the effect of ploidy on regulation of XCI can be further studied. HaESCs are established from *in vitro* cultured haploid embryos (Elling et al., 2011; Leeb and Wutz, 2011). HaESCs possess a haploid genome but also show a tendency towards diploidization, which is strongly enhanced when haESCs enter differentiation. Diploidization has, thus, far made studies of *Xist* activation difficult to interpret as haploid cells are in the process of diploidization. Several factors that affect diploidization have been extensively investigated in somatic lineage differentiation with the aim to reduce the high rates of diploidization (Freimann and Wutz, 2017; He et al., 2018; He et al., 2017; Olbrich et al., 2017; Olbrich et al., 2019; Takahashi et al., 2014). A previous study has reported the germline competence of haESCs *in vivo* by blastocyst injection (Leeb et al., 2012), but the ploidy of haESCs and their characterization during germ cell differentiation have been less well studied.

Here, we report the successful differentiation of haESCs into haploid PGCLCs *in vitro*. We show that a haploid genome is preferentially maintained in PGCLCs compared with somatic cell lineages. We then use this system to investigate *Xist* activation in haploid cells. Our data demonstrate that a single X chromosome is sufficient for *Xist* activation in a haploid genome consistent with a lower threshold of autosomal blocking factors.

## Results

### Differentiation of haESCs into haploid PGCLCs

To investigate the feasibility of germ cell differentiation from haESCs we followed a previously established protocol (Hayashi et al., 2012; Hayashi et al., 2011). A mixed population of haploid and diploid ESCs, containing 16.8% cells with a 1n DNA content (G0/G1/S-phase haploid), were differentiated to epiblast-like cells (EpiLCs) for 2 days in the presence of FGF2 and Activin A (Figure 1A, B). Subsequently, we aggregated the EpiLCs for embryoid body (EB) formation. PGCLCs appeared between day 6 and 8 and were identified by co-expression of the SSEA1 and integrin β3 surface markers. On day 7, 11.1% of the cells expressed both PGC markers (Figure 1C, D). Analysis of the ploidy distribution by flow cytometry after DNA staining with Hoechst 33342 showed that 9.3% of all cells possessed a 1n DNA content corresponding to haploid cells in G0-, G1- and S-phase, which is approximately half the fraction of haploid ESCs at the beginning of differentiation (16.8%; Figure 1E, F). To confirm a haploid karyotype in PGCLCs, we sorted cells expressing SSEA1 and integrin β3 from d7 EBs and prepared chromosome spreads. As expected, a set of 20 acrocentric chromosomes typical for an intact haploid mouse genome could be observed (Figure 1G). Transcription analysis of sorted PGCLCs revealed the upregulation of the PGC-markers *Blimp1, Prdm14, Tfap2c* and *Stella*, and the downregulation of *Dnmt3b* in G0/G1/S-phase haploid (1n) as well as in S/G2/M-phase diploid (4n) PGCLCs relative to EpiLCs and ESCs (Figure 2). These results show that differentiation of haploid PGCLCs *in vitro* recapitulates gene expression changes that are anticipated from the development of PGCs *in vivo*.

**Figure 1.**
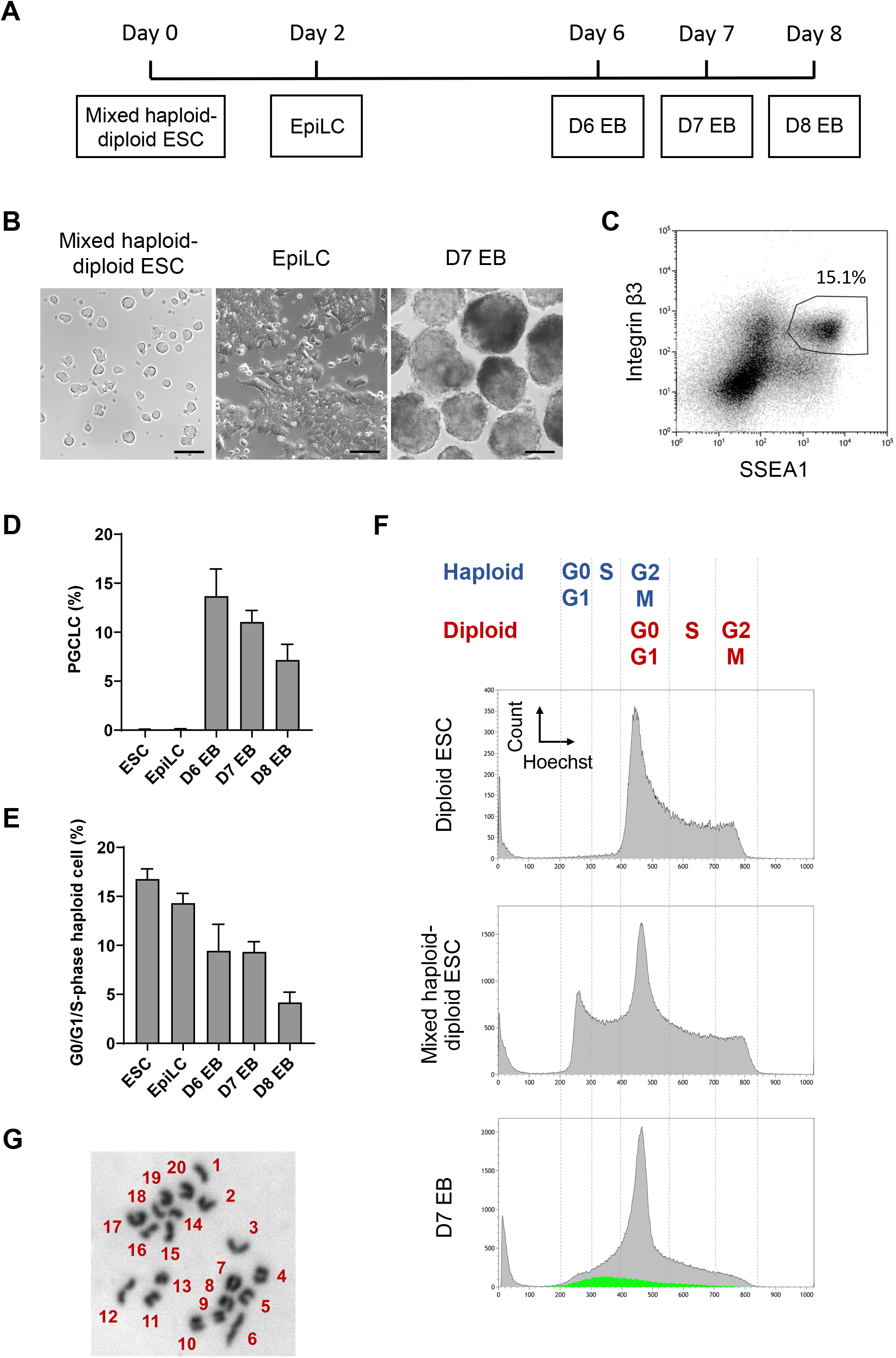
HaESCs differentiate to haploid PGCLCs by an in vitro culture. (A) A scheme of germ cell differentiation of mixed haploid-diploid ESCs. (B) Morphology of ESCs, EpiLCs and d7 EBs derived from mixed haploid-diploid ESCs. Scale bar, 100 µm. (C) A representative flow cytometry analysis of d7 EBs derived from mixed haploid-diploid ESCs. PGCLCs, positive for both SSEA1 and integrin β3, accounted for 15.1% out of all cells. (D, E) The proportion of PGCLCs (D) and G0/G1/S-phase haploid cells (E) out of all cells at the different stages during germ cell differentiation of mixed haploid-diploid ESCs. n = 14 (ESC), 9 (EpiLC), 7 (d6 EB), 18 (d7 EB) and 7 (d8 EB). Data represents the mean value and the standard error of the mean. (F) Flow cytometry analysis of DNA content of diploid ESCs, mixed haploid-diploid ESCs, and d7 EBs derived from mixed haploid-diploid ESCs. Cell cycle phases of haploid and diploid cells (top), and PGCLCs in d7 EBs (green) are indicated. (G) Representative chromosome spread of a haploid d7 PGCLC.

**Figure 2.**
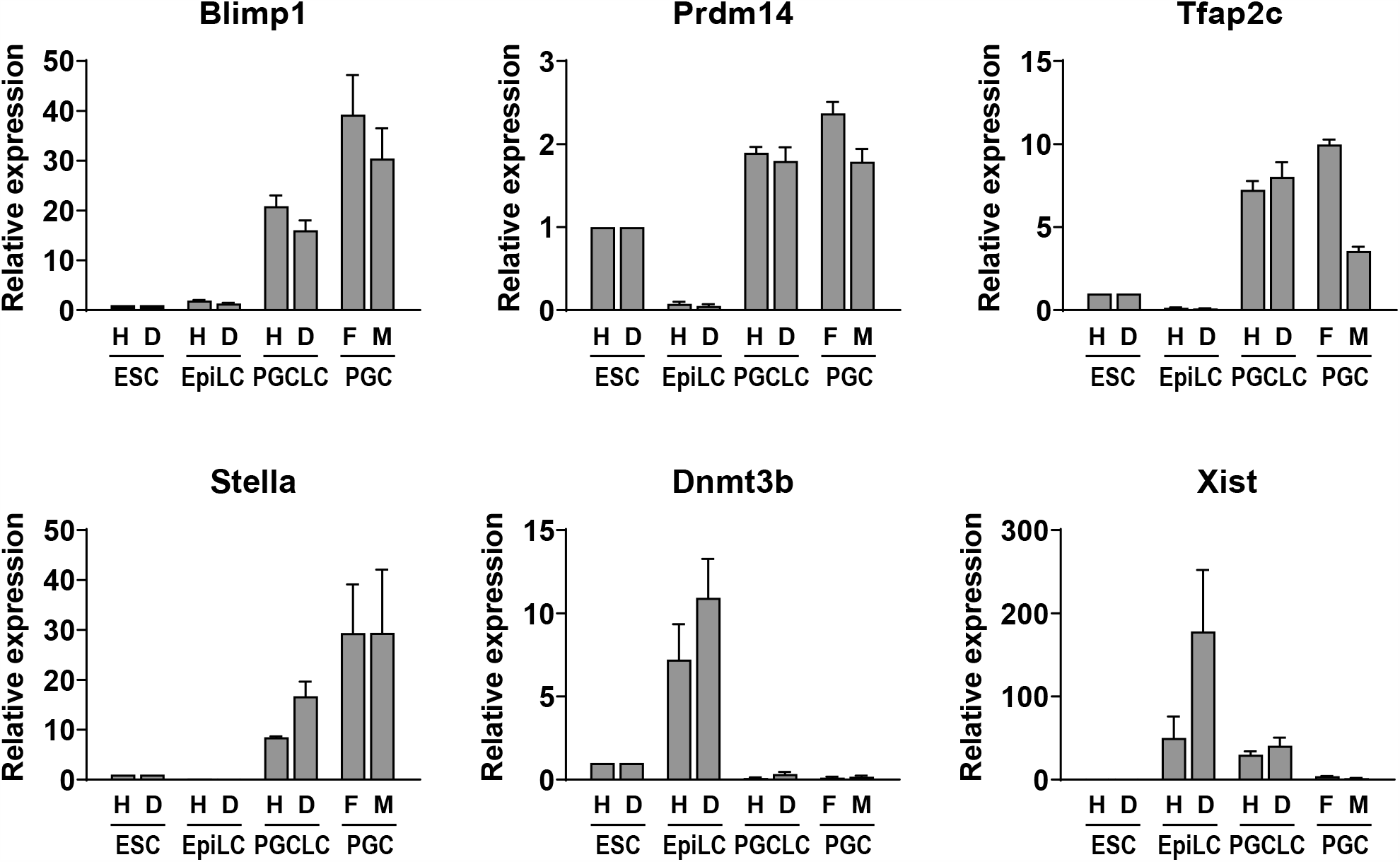
Transcription profile of haploid and diploid cells during germ cell differentiation *in vitro*. Transcription of PGC markers (*Blimp1, Prdm14, Tfap2c* and *Stella*), *Dnmt3b* and *Xist* during germ cell differentiation of mixed haploid-diploid ESCs. Gene expression of G0/G1/S-phase haploid and S/G2/M-phase diploid cells of ESCs, EpiLCs and d7 EBs was measured, respectively. Female and male PGCs purified from E12.5 gonads by FACS were used as controls. Gene expression of G0/G1/S-phase haploid and S/G2/M-phase diploid samples was normalized to *Gapdh* expression relative to G0/G1/S-phase haploid and S/G2/M-phase diploid ESCs, respectively. Gene expression of male and female PGCs was normalized to *Gapdh* expression relative to S/G2/M-phase diploid ESCs. Data represents relative expression of each sample with the mean value and the standard error of the mean (n = 4). H, G0/G1/S-phase haploid; D, S/G2/M-phase diploid; F, female; M, male.

### PGCLCs preferentially maintain a haploid genome

The PGCLC population of d7 EBs appeared to have a notably high content of haploid cells (Figure 3A, B). We therefore further charcteized the percentage of PGCLCs in windows of different DNA content. We analyzed the 1n population (G0/G1/S-phase haploid), the 2n population (G2/M-phase haploid and G0/G1-phase diploid), and 4n population (S/G2/M-phase diploid). In addition, we also analyzed the combined 2n and 4n population containing all but the 1n haploid cells. PGCLCs accounted for 32.3% of 1n (G0/G1/S-phase) haploid cells, while 11.1% of all cells and less than 10% of cell population with higher DNA content expressed both PGC markers (Figure 3C). Therefore, the 1n haploid population contained a 3-fold higher percentage of PGCLCs. The apparent enrichment of haploid cells within the PGCLC population can in part be attributed to high diploidization rates that accompany differentiation into somatic lineages. Nevertheless, a substantially higher percentage of haploid cells was observed in PGCLCs (27.5%) compared to overall population of ESCs (16.8%), EpiLCs (14.3%) and d7 EBs (9.3%; Figure 3D). These results indicate that PGCLCs preferentially maintain a haploid genome.

**Figure 3.**
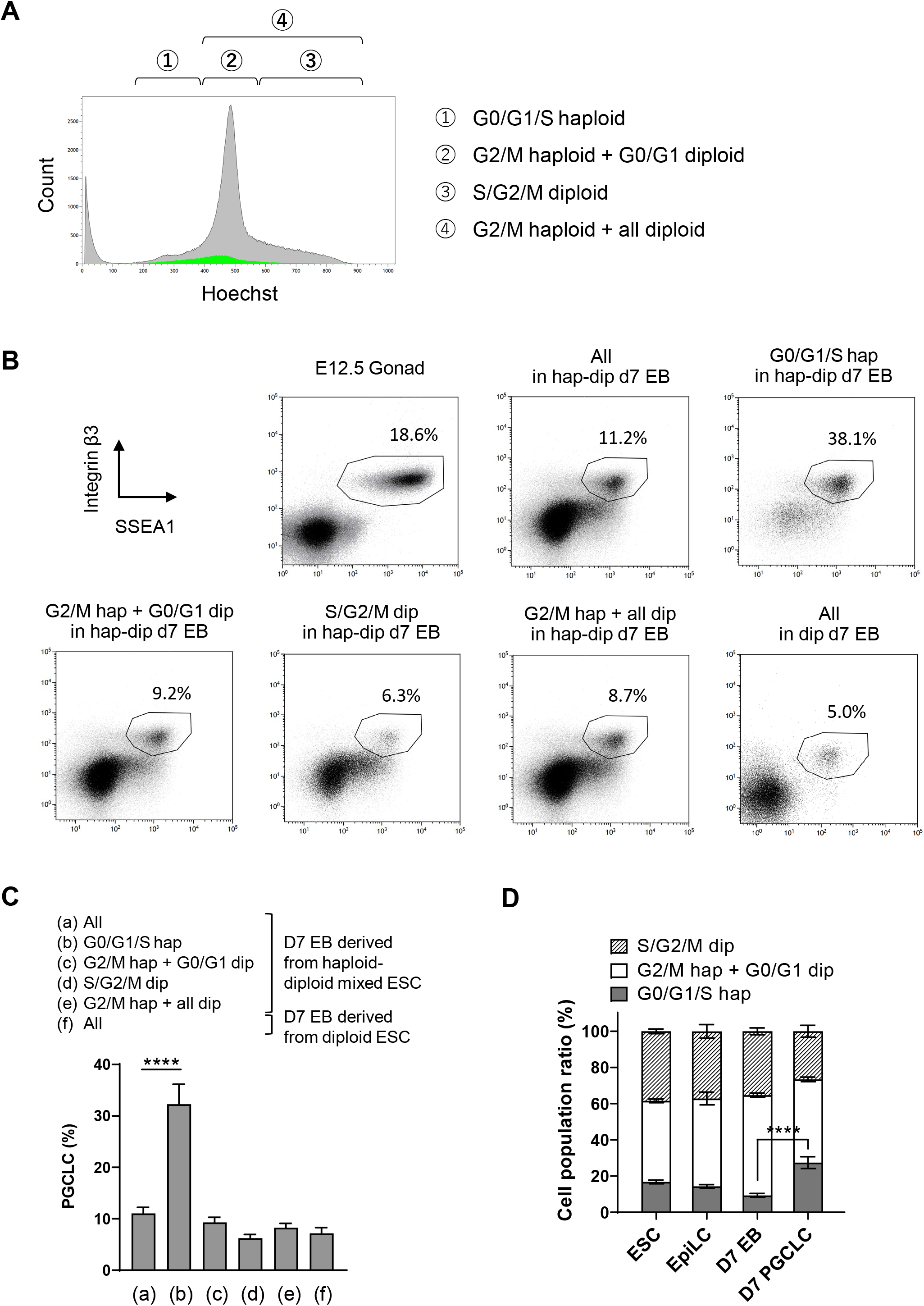
PGCLCs preferentially maintain haploidy *in vitro*. (A) A representative distribution of DNA content in d7 EBs derived from mixed haploid-diploid ESCs. PGCLCs, positive for SSEA1 and integrin β3, are indicated by green color. The cell cycle profile of haploid and diploid cells is shown at the top. (B) A representative flow cytometry analysis of d7 EBs. PGCLCs accounted for 11.2% out of all cells in d7 EBs derived from mixed haploid-diploid ESCs. PGCLCs accounted for 38.1%, 9.2%, 6.3% and 8.7% out of G0/G1/S haploid cell population, G2/M haploid + G0/G1 diploid cell population, S/G2/M diploid cell population, and G2/M haploid + all diploid cell population of d7 EBs derived from mixed haploid-diploid ESCs, respectively. Embryonic female gonads at E12.5 and d7 EBs derived from diploid ESCs were also analyzed as controls. (C) The ratio of PGCLCs in cell populations of different DNA content in d7 EBs derived from mixed haploid-diploid ESCs or diploid ESCs. PGCLCs accounted for 32.3% on average out of G0/G1/S haploid cells in d7 EBs derived from mixed haploid-diploid ESCs. Data represents the mean and the standard error of the mean. n = 18 (d7 EBs derived from mixed haploid-diploid ESCs) and n = 12 (d7 EBs derived from diploid ESCs). **** P < 0.0001. (D) Cell cycle distribution of haploid and diploid cells during germ cell differentiation of mixed haploid-diploid ESCs. Data represents the mean value and the standard error of the mean. n = 14 (ESC), 9 (EpiLC), 18 (d7 EB) and 18 (d7 PGCLC). **** P < 0.0001.

### *Xist* is activated from a single X chromosome in a haploid genome

Germline differentiation facilitates an investigation of *Xist* expression in haploid cells without caveats that arise from diploidization or cell death. We used sorted 1n (G0/G1/S-phase haploid) and 4n (S/G2/M-phase diploid) cells at different timepoints for *Xist* RNA FISH analysis. One and two punctate *Xist* signals were observed in haploid and diploid ESCs, respectively (Figure 4A). Our analysis aimed at the detection of *Xist* clusters and therefore used double stranded probes that also recognize nascent *Xist* and *Tsix* transcripts before initiation of XCI as a punctate focal signal. The majority of haploid and diploid ESCs showed no *Xist* cluster (Figure 4B). After the initiation of differentiation, *Xist* RNA clusters were observed in 59.0% of diploid EpiLCs as expected (Wutz and Jaenisch, 2000). A clear *Xist* cluster was also observed in 42.0% of sorted haploid EpiLCs indicating the initiation of XCI. *Xist* is activated from the single X chromosome in haploid cells with a similar frequency as from one of the two X chromosomes in diploid cells. Furthermore, 17.9% of diploid EpiLCs contained two separate *Xist* clusters. This observation might be explained by considering that the decision to activate *Xist* may have occurred before diploidization, when the cell was in a haploid state, and *Xist* expression was maintained on both X chromosomes after diploidization. Strong upregulation of *Xist* was also confirmed by quantitative RT-PCR analysis in both haploid and diploid EpiLCs (Figure 2).

**Figure 4.**
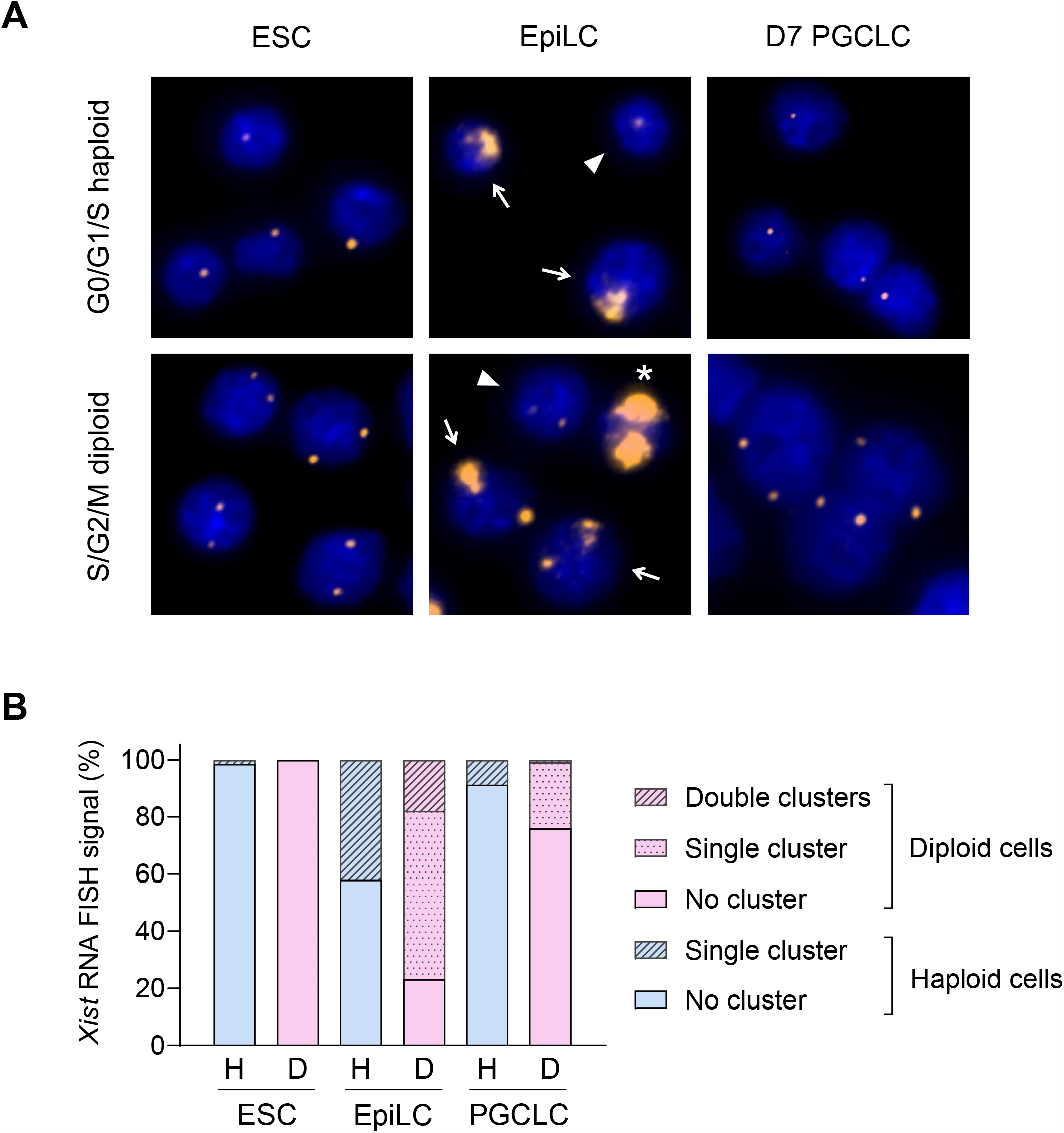
*Xist* is activated and repressed in both haploid and diploid cells during germ cell differentiation. (A) Representative images of *Xist* expression during germ cell differentiation of mixed haploid-diploid ESCs detected by RNA FISH using a Cy3-labeled *Xist* probe (orange). No *Xist* cluster but single foci observed in G0/G1/S-phase haploid ESCs and PGCLCs. No cluster but double foci observed in S/G2/M-phase diploid ESCs and PGCLCs. Single *Xist* clusters (arrow) and foci (arrowhead) observed in G0/G1/S-phase haploid EpiLCs. Double clusters (asterisk), single clusters (arrow) and no cluster (arrowhead) observed in each S/G2/M-phase diploid EpiLC. (B) Proportion of *Xist* RNA FISH signals during germ cell differentiation of mixed haploid-diploid ESCs. The number of cells possessing no, single or double *Xist* RNA clusters were counted in G0/G1/S-phase haploid and S/G2/M-diploid cells. Total numbers of counted cells are 138 (H, ESC), 171 (D, ESC), 100 (H, EpiLC), 173 (D, EpiLC), 69 (H, PGCLC) and 121 (D, PGCLC). D, S/G2/M-phase diploid; H, G0/G1/S-phase haploid.

With further differentiation, we found that the proportion of cells exhibiting *Xist* clusters decreased (Figure 4A, B). Only 8.7% and 24.0% of haploid and diploid d7 PGCLCs showed *Xist* clusters, respectively. This observation might reflect the repression of *Xist* in germ cell differentiation and cell selection that is expected from the loss of X-linked gene expression in haploid cells after inactivation of the single X chromosome. The ability to repress *Xist* may contribute to the ability of the germline to maintain a haploid genome with a higher frequency than somatic lineages that normally do not express repressors of *Xist*. Taken together, our data demonstrate that *Xist* is activated from a single X chromosome without *Xic* pairing in differentiating haESCs.

## Discussion

Our observation that haESCs maintain a haploid genome during germ cell differentiation enabled us to analyze *Xist* activation in the context of a haploid genome. Diploid mouse ESCs activate *Xist* upon entering differentiation and have been used as a model for studying the underlying regulation of XCI. Our results are explained by the idea that the amount of blocking factors produced from a single set of autosomes is insufficient to counteract activators from a single X chromosome. In contrast, X-linked activators are titrated by a double dose of blocking factors in diploid male cells preventing *Xist* activation. Our result therefore supports a model of diffusible X-linked activators and autosomal blocking factors (Barakat et al., 2014; Pollex and Heard, 2019). We note that haploid cells have a smaller cell volume than diploid cells (Freimann and Wutz, 2017), and it is not entirely clear if cell volume could have an influence on XCI. Previous studies have also linked the activation of *Xist* with *Xic* pairing in differentiating diploid ESCs (Bacher et al., 2006; Xu et al., 2006). *Xic* pairing cannot occur in haploid cells as only a single X chromosome is present. Our experiment shows that *Xist* activation does not strictly depend on *Xic* pairing. This observation does not rule out that *Xic* pairing contributes to XCI in diploid cells or has a role in ensuring that one X chromosome remains active after the decision for initiating XCI has been taken in female cells. A recent study has reported on engineering the *Xic* regions for tethering to the nuclear lamina (Pollex and Heard, 2019). XCI was initiated in female mouse ESCs despite *Xic* movement was restricted.

Previously, *XIST* expression has been analyzed in human haESCs (Sagi et al., 2016). XCI in the human embryo is regulated differently from the mouse. *XIST* is initially expressed from the single and both X chromosomes in male and female embryos, respectively (Okamoto et al., 2011). In contrast to mouse, *XIST* does not lead to chromosomal silencing in human preimplantation embryos. The *XACT* gene, which is not present in mice, prevents silencing and a partial repression of X-linked genes is observed that has been termed “dampening” (Sahakyan et al., 2017). After implantation *XIST* expression becomes monoallelic in female embryos and XCI is initiated. Conversely, the post implantation human male embryo ceases *XIST* expression. Female human ESCs show monoallelic *XIST* expression and an Xi. During culture erosion of the Xi has been observed, which is characterized by a loss of *XIST* expression and reactivation of genes from the Xi to various extent. Established human haESC lines have been shown to lack *XIST* expression (Sagi et al., 2016). Although, this observation is expected, it is not entirely clear how an *XIST* negative state was established. Lack of *XIST* activation, selection of cells with an active X chromosome, and the erosion of an Xi can be considered. The fact that *XIST* is initially expressed from all X chromosomes in human diploid preimplantation embryos, suggests that it is also activated in haploid embryos. It is conceivable that mechanisms that prevent inactivation including *XACT* could contribute to survival and potentially lead to repression of *XIST* in human haESCs (Vallot et al., 2017). Irrespectively, it is likely that the lack of *XIST* expression in established haESC lines does not reflect the state of the embryonic cell of origin. In contrast, mouse ESCs have not initiated XCI. Capturing of the earliest stages of XCI upon entry into differentiation permits the study of the X chromosome counting mechanism in culture.

Our study also reveals that PGCLCs maintain a haploid genome with high frequency. Therefore, the diploidization rate during germline differentiation is comparable to haESCs and substantially lower than that of somatic lineages. PGCs have a similar epigenetic state to ESCs, which might contribute to tolerance of a haploid genome. Firstly, during migration to the gonads, the Xi becomes reactivated suggesting that dosage compensation is not essential for germ cell development. Secondly, ESCs and PGCs exhibit genome-wide DNA hypomethylation (Leitch et al., 2013). Lastly, transcription factors that are associated with pluripotent cells and germ cells including Oct4 have been implicated as repressors of *Xist* (Donohoe et al., 2009; Navarro et al., 2010; Nesterova et al., 2011). It is conceivable that the expression of potential repressors of *Xist* in PGCs might contribute to the maintenance of a haploid genome consistent with our finding that *Xist* expression is lost at later time points in differentiating cultures. Oct4 is also expressed in EpiLCs and, thus, cannot explain repression of *Xist* in ESCs and PGCs. However, other factors have also been implicated in *Xist* repression in pluripotent cells. Haploid PGCLCs will be useful for future studies of ploidy restriction and the genetics of germ cell development.

## Experimental Procedures

### Derivation and culture of haESC lines

Derivation of haESC lines from 129S6/SvEvTac mice was performed as previously described (Leeb and Wutz, 2011). At passage 5, G0/G1/S-phase haploid cell population of a haESC line was purified by fluorescence-activated cell sorting (FACS) using flow cytometer (MoFlo Astrios EQ, Beckman Coulter) after Hoechst 33342 (Invitrogen) staining. Sorted cells were cultured and maintained on a gelatin-coated plate with irradiated mouse embryonic fibroblasts (MEFs) of DR4 mouse embryos at E12.5 (The Jackson Laboratory, no. 003208). Cells were maintained in Serum+2i+LIF medium, which was mixed with equal volume of Serum+LIF medium without 2i (Postlmayr et al., 2020) and 2i+LIF medium (Hayashi and Saitou, 2013). At passage 10, G0/G1/S-phase haploid cell population of the haESC line was purified by FACS and was maintained on a gelatin-coated plate with MEFs in Serum+2i+LIF medium.

### *In vitro* germ cell differentiation

Germ cell differentiation was performed following a published protocol (Hayashi and Saitou, 2013) with a few modification. The haESC line was cultured on an ornithine- and laminin-coated plate without MEFs in 2i+LIF medium from passage 12. At passage 15, EpiLC differentiation was initiated as described in the protocol. After 2 days of EpiLC differentiation, 2.3 × 10^5^ EpiLCs were plated into a well of a Sphericalplate 5D (Kugelmeiers Ltd.) with 1.4 ml of PGCLC differentiation medium not containing BMP8a. After 4 days of PGCLC differentiation, half of the medium was replaced with fresh PGCLC differentiation medium not containing BMP4 and BMP8a.

### Flow cytometry analysis and cell sorting

To investigate the cell cycle of haploid and diploid cells and PGCLCs, flow cytometry analysis was performed to ESCs, EpiLCs and EBs by following procedures. Cells were harvested from culture vessels as described in a published protocol (Hayashi and Saitou, 2013), followed by staining with 15 µg/ml Hoechst 33342 for 12 min at 37 °C. Subsequently, PE anti-integrin β3 (BioLegend, no. 104307) and eFluor 660 anti-SSEA1 (eBioscience, no. 50881341) were added to the cell suspension at concentration of 1 µg/ml and 0.12 µg/ml, respectively, and the cell sample was kept on ice for 12 min. The fluorescence of each dye was measured by flow cytometry. Cell cycle of haploid and diploid cells was determined based on peaks of cell population at 1n and 2n DNA contents, corresponding to G0/G1-phase haploid cells and G2/M-phase haploid and G0/G1-phase diploid cells, respectively, by measuring the signal of Hoechst 33342. The population of PGCLCs was determined by measuring the signals of PE and eFluor 660.

### Karyotyping

On day 7 of germ cell differentiation, M-phase arrest of EBs was performed by culturing in PGCLC differentiation medium supplemented with 0.05 mg/ml demecolcine (Merck) for 8 hours. Subsequently, PGCLCs were sorted by FACS. Chromosome counting of PGCLCs was performed as previously described (Aizawa et al., 2020).

### Transcription analysis

During germ cell differentiation of mixed haploid-diploid ESCs, G0/G1/S-phase haploid and S/G2/M-phase diploid ESCs, EpiLCs and d7 PGCLCs were sorted from ESCs, EpiLCs and d7 EBs by FACS, respectively. E12.5 gonads were harvested from 129S6/SvEvTac mouse embryos. Sex of the gonads was determined by their morphology. PGCs were identified as gonadal cells positive for both SSEA1 and integrin β3 and were sorted by FACS. RNA of each sample was extracted using the RNeasy Mini Kit (Qiagen) following the manufacturer’s protocol, including an on-column DNA digest using RNase-free DNase (Qiagen). RNA concentration was determined using a NanoDrop Lite (Thermo Fisher Scientific). 100 − 500 ng total RNA was reverse transcribed using the PrimeScript RT Master Mix (Takara) according to the manufacturer’s instruction. RT-PCR was performed on a 384 well format with the 480 Lightcycler instrument (Roche) using KAPA SYBR FAST qPCR KIT (Kapa Biosystems). Fold change expression was calculated using the ΔΔct method. *Gapdh* expression of G0/G1/S-phase haploid ESCs was used to normalize the transcription of G0/G1/S-phase EpiLCs and d7 PGCLCs. *Gapdh* expression of S/G2/M-phase diploid ESCs was used to normalize the transcription of S/G2/M-phase EpiLCs, d7 PGCLCs and PGCs. Primers used for transcription analysis are listed in Table S1.

### *Xist* RNA FISH

RNA FISH was performed to analyze *Xist* expression during germ cell differentiation of mixed haploid-diploid ESCs. FISH probe was prepared from ptetOP-Xist-PA plasmid with Cy3-dCTP (Amersham Biosciences) as described previously (Wutz and Jaenisch, 2000). G0/G1/S-phase haploid and S/G2/M-phase diploid ESCs, EpiLCs and d7 PGCLCs were sorted by FACS, respectively. Sorted cells were mounted onto glass slides using Cytospin 4 (Thermo Scientific) for 3 min at 800 or 900 rpm for haploid or diploid cells, respectively.

Glass slides were immediately rinsed in PBS and washed in CSK buffer (100 nM NaCl, 300 nM sucrose, 3 mM MgCl_2_, 10 mM PIPES pH 6.8) for 30 sec, in CSK buffer + 0.5% Triton X-100 for 2 min and again in CSK buffer for 30 sec. Cells were fixed in 4% paraformaldehyde in PBS for 10 min and washed in 70%, 80%, 95% and 100% EtOH for 2 min each. After airdrying the slides, hybridization was performed by applying probe to the cells. Probes were covered with a coverslip and sealed with rubber cement. Slides were then placed in a humidified chamber and incubated overnight at 37°C. Coverslips were removed and slides were washed in 2X SSC + formamide (50%) for 15 min at 39°C, in 2X SCC three times for 5 min each at 39°C, in 1X SSC for 10 min and in 4X SSC with a dip (20X SSC: 3 M NaCl, 0.3 M tri-sodium citrate dihydrate in H_2_O). Cellular DNA was counterstained by incubating slides in 4X SSC + 0.1% Tween + 14 mM DAPI for 1.5 min. Slides were washed with 4X SSC for 5 min. After washing, the sample was mounted in Vectashield (Vector Laboratories) and covered with a coverslip. Samples were imaged under the microscope (Axio Observer Z1, Zeiss) equipped with an ORCA-Flash4.0 camera (Hamamatsu Photonics K.K.). Images were processed using Zeiss Zen Pro 2.0 software.

### Statistical analysis

For comparison of cell population ratio (Figure 3C, D), measurements were analyzed with the GraphPad Prism 8 software using a two-tailed t-test. A p-value < 0.05 was considered statistically significant.

## Acknowledgments

We thank Dr. Giulio Di Minin for derivation of haESC lines. This study was supported by the Swiss National Science Foundation (grant 31003A_152814/1).

## Author Contributions

E.A. and A.W. conceptualized experiments. E.A., C.K., S.S. and R.F. collected the data.

E.A. and C.K. analyzed the data. E.A. and A.W. wrote the manuscript.

## Supplemental Information

**Table S1.**
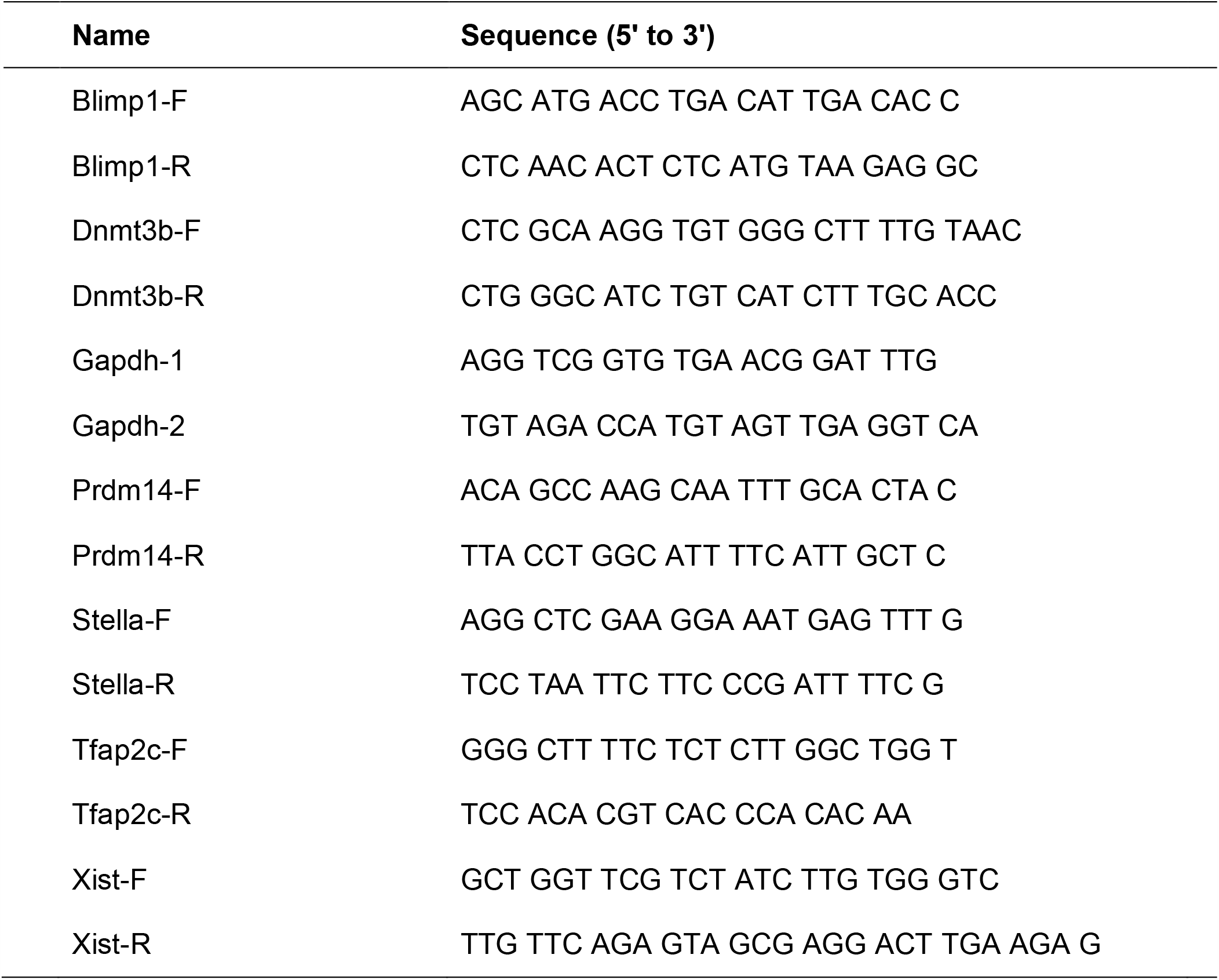
List of oligos

