## Supplementary figures and images for "Haploid mouse germ cell precursors from embryonic stem cells reveal *Xist* activation from a single X chromosome" (from another version of this preprint: version 1, 2020-10-30)

### Figure S1

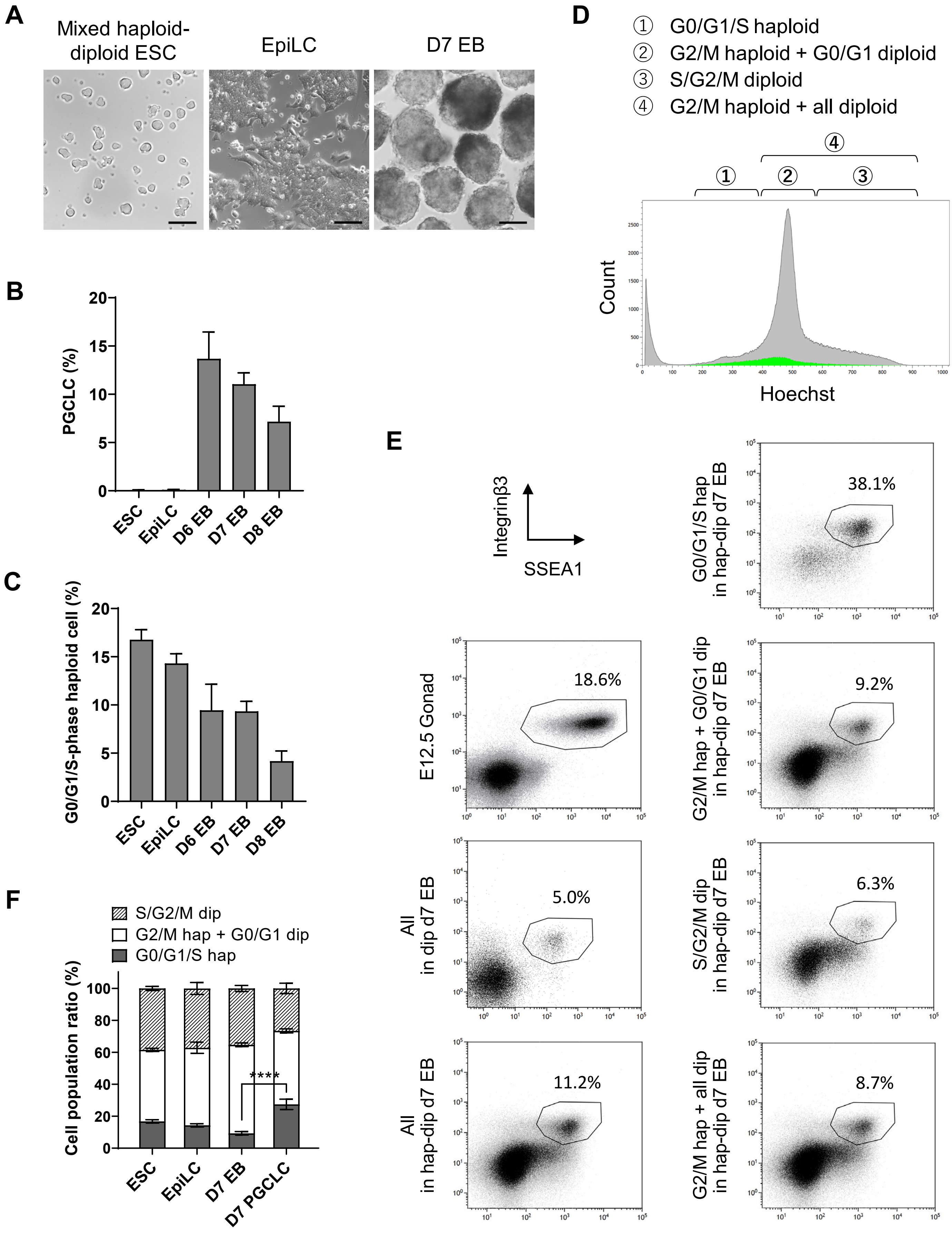

### Figure S2

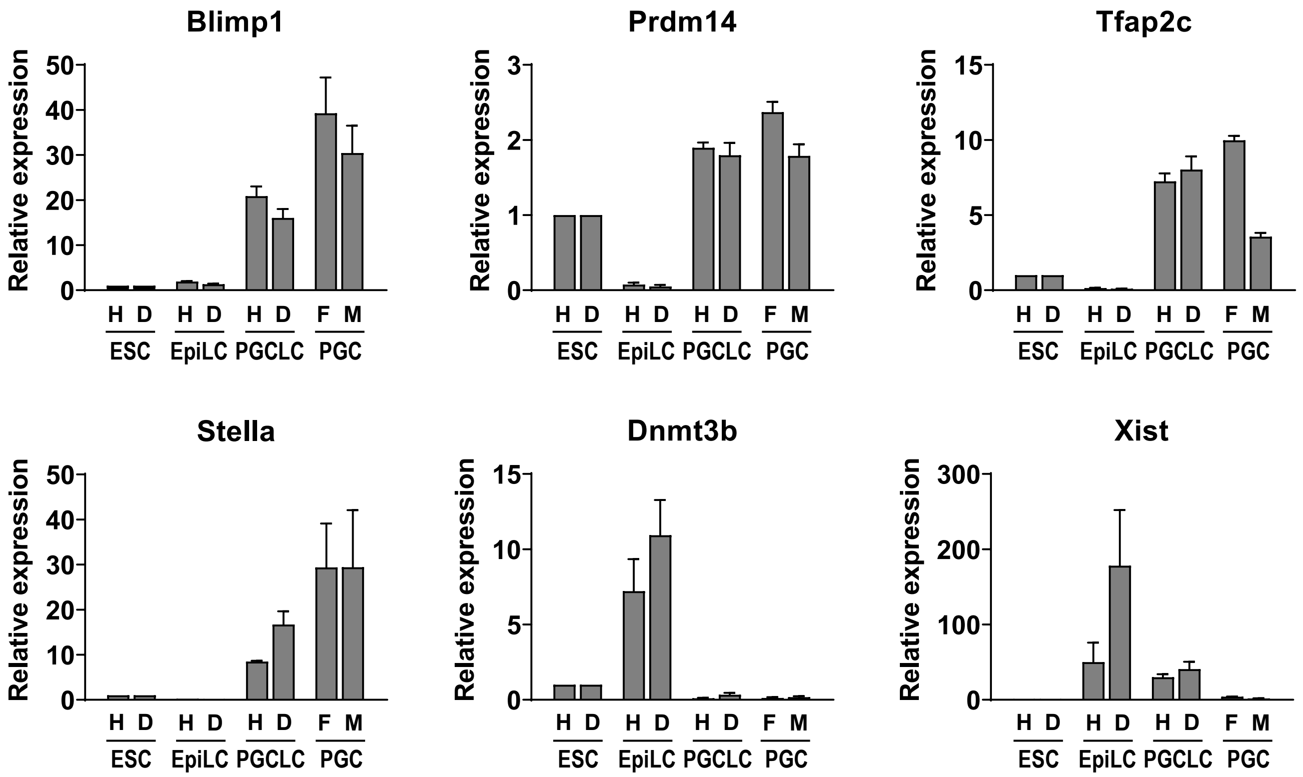

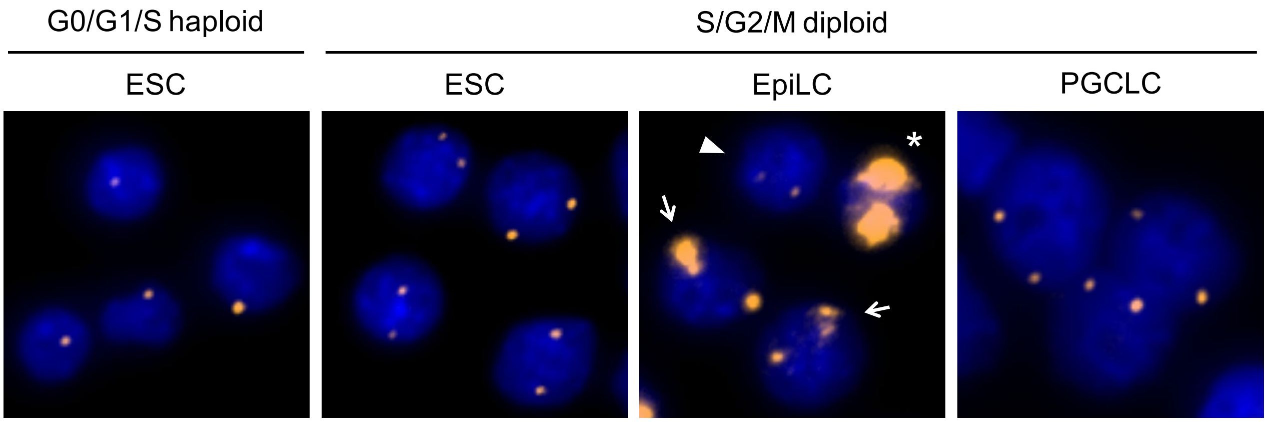
