## Supplementary material for "Haploid mouse germ cell precursors from embryonic stem cells reveal *Xist* activation from a single X chromosome": Table S1 (from another version of this preprint: version 1, 2020-10-30)

### Table S1. List of oligos

| Name | Sequence (5' to 3') |
|---|---|
| Blimp1-F | AGC ATG ACC TGA CAT TGA CAC C |
| Blimp1-R | CTC AAC ACT CTC ATG TAA GAG GC |
| Dnmt3b-F | CTC GCA AGG TGT GGG CTT TTG TAAC |
| Dnmt3b-R | CTG GGC ATC TGT CAT CTT TGC ACC |
| Gapdh-1 | AGG TCG GTG TGA ACG GAT TTG |
| Gapdh-2 | TGT AGA CCA TGT AGT TGA GGT CA |
| Prdm14-F | ACA GCC AAG CAA TTT GCA CTA C |
| Prdm14-R | TTA CCT GGC ATT TTC ATT GCT C |
| Stella-F | AGG CTC GAA GGA AAT GAG TTT G |
| Stella-R | TCC TAA TTC TTC CCG ATT TTC G |
| Tfap2c-F | GGG CTT TTC TCT CTT GGC TGG T |
| Tfap2c-R | TCC ACA CGT CAC CCA CAC AA |
| Xist-F | GCT GGT TCG TCT ATC TTG TGG GTC |
| Xist-R | TTG TTC AGA GTA GCG AGG ACT TGA AGA G |
